# RedMed: Extending drug lexicons for social media applications

**DOI:** 10.1101/663625

**Authors:** Adam Lavertu, Russ B Altman

## Abstract

Social media has been identified as a promising potential source of information for pharmacovigilance. The adoption of social media data has been hindered by the massive and noisy nature of the data. Initial attempts to use social media data have relied on exact text matches to drugs of interest, and therefore suffer from the gap between formal drug lexicons and the informal nature of social media. The Reddit comment archive represents an ideal corpus for bridging this gap. We trained a word embedding model, RedMed, to facilitate the identification and retrieval of health entities from Reddit data. We compare the performance of our model trained on a consumer-generated corpus against publicly available models trained on expert-generated corpora. Our automated classification pipeline achieves an accuracy of 0.88 and a specificity of >0.9 across four different term classes. Of all drug mentions, an average of 79% (±0.5%) were exact matches to a generic or trademark drug name, 14% (±0.5%) were misspellings, 6.4% (±0.3%) were synonyms, and 0.13% (±0.05%) were pill marks. We find that our system captures an additional 20% of mentions; these would have been missed by approaches that rely solely on exact string matches. We provide a lexicon of misspellings and synonyms for 2,978 drugs and a word embedding model trained on a health-oriented subset of Reddit.

## 1. Introduction

Social media represents a key opportunity for gathering real-world evidence regarding drug safety and efficacy. The U.S. Food and Drug Administration (FDA) and others have recognized the utility of social media and have called for further research [1, 2]. Indeed, social media is already in use by several drug monitoring systems, such as RADARS and NDEWS [3, 4]. Social media has a large and growing user base within the United States with 69% of adults reporting the use of at least one social media platform [5]. On an average day, users generate more than 2.5 million Reddit comments and more than 500 million Tweets.

The result of this large amount of activity is a massive high-frequency data stream that has the potential to revolutionize the way we study drugs. Social media offers access to an observational data source that is potentially orthogonal to data streams that are currently being utilized for gathering real-world data, such as spontaneous event reporting systems, electronic health records, and surveys. The pseudo-anonymous nature of some of these social media platforms might make users more inclined to discuss stigmatized behaviors of potential interest to researchers [6].

It is currently difficult to identify social media content regarding drugs at scale. Given the massive size of these data sets, a first step in most analyses is the identification of relevant user activity. This first step is crucial and often performed via APIs that are necessary to access the social media data. These APIs generally require explicit search terms to identify the data to be returned to the user. Once data is successfully retrieved, it can be processed by context aware named entity recognition models for downstream research applications. Initial efforts have generally relied on identifying relevant social media content through exact lexical matching to known generic and trademark drug names [7, 8, 9, 10, 11]. Throughout the rest of this paper, we refer to these exact lexical matches as “known” terms. A few analyses have extended exact match lexicons through the manual curation of ad-hoc single-use lists of search terms for their drugs of interest [12, 13, 14, 15]. Unfortunately, due to the informal nature of social media text derived from these platforms displays greater variability than texts from more formal sources, such as Wikipedia or PubMed [16]. The informality of the medium very likely results in many drug mentions being missed, such as those that are slang terms for and/or misspellings of the drug of interest. A recent small scale study, found around 10% of drug mentions on Twitter were misspelled, with higher rates of misspellings for drugs of abuse [15].

A few efforts have focused specifically on the task of generating lists of potential misspellings of drug names. Work by Pimpalkhute et al. was one of the first efforts to attempt to algorithmically generate these potential misspellings [17]. Their approach first generated a list of all potential misspellings that were within 1-edit distance of a given drug name, then filtered those candidates based on phonetic similarity between the drug name and the proposed misspelling. The final list of candidates was passed through an additional filter based on the number of search Google search results returned for a query of the proposed misspelling plus the word “drug”. Their use of a maximum of 1-edit distance from the original drug name severely limited their effort but was done to avoid the combinatoric explosion that results from allowing multiple edits. They made no use of semantic information in their approach, however their use of Google search results as a means of validation seems to have been effective and is expanded upon in our effort. Sarker et al. made additional progress towards the goal of generating potential misspellings. [18] This effort is notable for it’s combination of both semantic and lexical information, as well as it’s use of a recursive search strategy. They utilized a word embedding model to identify tokens that are semantically similar to a given query term and then filtered the resulting list using weighted Levenshtein ratio, a measure of lexical similarity. The cutoff for the lexical filter was tuned using a development set of labeled misspellings. They are similar to our effort in their use of both semantic and lexical information, but differ in their use of manually curated labels for tuning a cutoff threshold and their consideration of a single drug at a time, rather than a collection of known drug names. So a search misspellings of “atorvastatin” could potentially result in a false positive that is another known drug, such as “rosuvastatin”. While the research effort from which they derived their word embeddings explored several parameter settings, it did not optimize for any particular characteristics within the embedded space [19]. Additional work has been done to identify keywords consisting of both misspellings and slang terms. Simpson et al. performed a study to evaluate the validity of word embeddings models train on social media data for the task of identifying these misspelling and slang terms [13]. They first trained a continuous bag of words (CBOW) model on 86.2 million Tweets collected from Twitter in July of 2016. They used a set of marijuana seed terms to produce a list of candidate terms based on cosine similarity in the embedded space and selected a cutoff using a set of crowd-sourced marijuana slang terms. They manually curated the final list of 200 candidate terms and found 32.5% of the candidate terms to be direct references to marijuana. Their method has limited scalability due to the need for human list curated list of true positives against which to determine a cutoff for each drug, as well as the reliance on human evaluation of the final candidate term list. Adams et al. extended this work to consider the interplay between a particular keyword identification method and the corpus to which that method was applied [14]. They trained separate CBOW word embedding models on Twitter and Reddit corpora, then evaluated the effectiveness of two different search strategies to identify misspellings and phrases in the respective vector spaces. Their analysis was limited to identifying terms related to marijuana or opioids and their search strategy used only semantic information from the embedded space. Notably, they found a Reddit corpus filtered for subreddits related to marijuana and opioids greatly increased the performance of their keyword identification method. Therefore, a more comprehensive set of social media lexicons is a critical missing element for the identification of social media data of interest. Our approach incorporates elements of the approaches used in these efforts and addresses several of their associated limitations.

Word embedding models can capture semantics and are an ideal method for the discovery of misspellings and synonym terms [20]. These models rely on the distributional hypothesis, that the meaning of a word is represented by the meaning of the words it co-occurs with in context [21]. This hypothesis is represented by word co-occurrence statistics, where a model learns weights while being trained to predict which words are likely to co-occur and which are not. The learned model weights can then be used as vectorized representations of the words themselves. Previous work has demonstrated that these vectorized representations effectively represent word meanings and can be used in combination with vector operators to facilitate different language tasks [20]. Using word embeddings to represent terms within a social media corpus, we may be able to identify potential misspellings and synonyms of given term based on vector distance.

The Reddit comment archive (RCA) represents a large publicly available corpus of text generated in a social media setting. Reddit is the 5th most visited website in the United States, has more than 330 million active users, and an average of 21 billion screen views per month [22, 23]. A primary feature of the Reddit social media platform is the existence of subreddits, moderated discussion boards with a focused topic. Subreddit discussions span a broad range of topic areas, from politics to illicit drug use to health conditions. We focused on a subset of subreddits related to health, as determined by the number of mentions of generic/trademark drugs and adverse drug reactions (ADRs) contained within the subreddits. A few examples of these health oriented subreddits are ‘r/trees’, for the discussion of marijuana, ‘r/opiates’, for the discussion of opiates, ‘r/diabetes’, discussion of diabetes and diabetes management, and ‘r/fitness’ focusing on general fitness. A single discussion thread consists of an initiating user submission called a post and associated threads of user comments responding to both the initial post and to other comments.

The RCA data has several advantages for the purpose of identifying drug synonyms and misspellings compared to other previously used corpora. First, the RCA contains comments created by individuals with little or no editing (i.e. unlike Wikipedia), thus it preserves both spelling errors and potentially rarer semantic usage of community specific terms, such as slang. Second, the subreddit structure of Reddit, as mentioned above, offers high-level metadata about the likely topic of a given comment in a particular subreddit. This is comparable to the usage of hashtags on Twitter to denote a topic of relevance, but unlike hashtags which aren’t required for every individual tweet, all Reddit comments are annotated with their subreddit of origin. A recent study found the ability to filter Reddit content based on subreddits improved the performance of their method over the use of unfiltered Twitter data [14]. However, if metadata about users, such as age, sex, or location, is of great interest to a research effort, Twitter user profiles often contain such metadata at higher rates than Reddit. Third, the language in the RCA is informal and yields a training corpus that is representative of average language usage on social media. Fourth, Reddit comments are conversational and contain many individual mentions of a particular terms in context. Word embedding models, which rely on the distributional hypothesis, benefit from having many different contextualized examples of terms. Fifth, while Reddit has fewer users and daily post volumes than Twitter, Reddit only places a 10,000-character limit on comments/posts. In contrast, Twitter, limited posts to 140 characters until November 2017 and currently limits them to 280 characters [24]. Sixth, the RCA is publicly archived and fully downloadable without restrictions on data storage and sharing. This facilitates research reproducibility as authors can refer to the archived comment data. Although, Twitter data can be fully archived during the original research effort and shared with other researchers upon request. Recent independent efforts to replicate studies that used Twitter data were only able to retrieve 1,012 of the 1,784 tweets included in the original 2015 study, 57% of the original data set [25].

Our work uses the RCA, word embedding models, and automated filters to create an empirical lexicon of potential drug misspellings and synonyms. We trained a word embedding model, RedMed, on Reddit data and optimized it for the retrieval of drug synonyms. We used a database of known drug terms and a novel performance metric, as well as a novel word cluster pruning technique in conjunction with the model to identify candidate drug terms. We then filtered these candidate tokens using lexical edit-distance, phonetics, pill marks, and Google search results. We performed manual validation on a subset of these results and achieved an accuracy of ~0.88. The utility of this lexicon is demonstrated by increased yield of relevant user posts. These drug term lists represent a novel resource for efforts in social media-oriented drug research.

## 2. Materials and methods

### 2.1. Dataset

#### 2.1.1. Reddit comment data and subreddit selection

Reddit comments from 2005-2018 and their associated metadata were downloaded from https://pushshift.io [26]. In order to maximize the ability of models to learn health-oriented semantics from the corpus, we sought to create subsets of the corpora that were enriched for health-related content by leveraging comment metadata with regards to what subreddit each comment was submitted to. This allows for the broad selection of comments that were made within subreddits dedicated to health-related topics. We selected an initial training corpus for the word embedding models by extracting all forums listed in the r/Health and r/Drugs side bars. After an initial hyperparameter optimization utilizing this corpus, we searched all comments in the RCA, regardless of the subreddit, for exact matches to generic and trademark drugs contained within DrugBank and health-related terms derived from the SIDER subset of MedDRA [27, 14, 28, 29]. This data-driven approach enabled the inclusion of a greater number of health-related subreddits that were not curated within the side bars mentioned above, such as subreddits that were recently created, were banned but archived, contained low numbers of subscribers, and/or are dedicated to more obscure health topics. We then selected the final collection of subreddits to include based on their enrichment for drug mentions per comment (i.e. number of comments within a subreddit containing a drug mention divided by the total number of comments within that subreddit). We selected an enrichment cutoff of 1*e*^−4^, which was the point at which the number of unique drugs captured within the corpus started to plateau.

#### 2.1.2. de novo phrase finding

The lack of a comprehensive list of consumer drug phrases necessitated the *de novo* discovery of phrases (e.g. word level n-grams that are enriched within the corpus). We utilized AutoPhrase [30, 31], a phrase discovery tool, to identify phrases within the initial development corpus derived from the r/Health and r/Drugs side bars. AutoPhrase requires a set of training phrases to calibrate the scoring scheme, we used 70,108 phrases contained in Wikipedia, MeSH, DrugBank, and the Consumer Health Vocabulary [32] as the training data. AutoPhrase was run without dependency parsing and phrases with a score above 0.5 were utilized for corpus processing. At this stage phrases were not filtered for relevance to health or drugs, but 2,849 of these phrases contained the generic name of a drug from DrugBank.

#### 2.1.3. Corpus preprocessing

Each comment in the corpus was preprocessed using the following steps, (1) all text was normalized to lowercase, (2) text segments corresponding to a discovered phrase were joined into a single token, (3) URLs and links to other users and subreddits were removed, (4) all tokens were split on non-word characters unless they were a phrase from 2.1.2, (5) stop words were filtered from the remaining token sets (Supp. File 1). Duplicate comments were removed.

### 2.2. Word embedding model

#### 2.2.1. Model training

A continuous bag-of-words model [20] was trained using Python 3.5 and the gensim package v2.3 [33]. We used an embedding size of 64 dimensions, a window size of 7 and a minimum count of 5. A hyperparameter optimization of the model was performed through a grid search over the negative sampling exponent parameter from −1.0 to 1.0 by steps of 0.25 [34]. This resulted in the selection of 0.25 as the negative sampling exponent (default value is 0.75). All remaining hyperparameters were set to the defaults. The model was trained for 25 iterations, at which point the performance on the DrugBank Clustering Coefficient task, detailed below, had plateaued.

#### 2.2.2. DBCC: DrugBank Clustering Coefficient

We used DrugBank to create a similarity task for word embedding model evaluation. Trademark and generic drug names were grouped based on their DrugBank identifier. The DBCC was then used to measure the relative similarity of grouped drugs versus drugs not in the same DrugBank group.

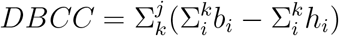

Where *j* is the set of all drug groups and *k* is the set of drugs contained within an individual drug group. We then sum *b*_*i*_, the pairwise cosine similarities for all drugs within that drug group, and subtract *h*_*i*_, the cosine similarity of a random drug within the set and a random drug outside the set, for an equivalent number of drug pairs to the drug pairs within the group. The sum of the difference in pairwise similarity across all drug groups is the DBCC score for that model. The DBCC has a high value when the vectors for trademark and generic names for a particular compound have a higher cosine similarity within the embedded space than random drug pairs, indicating that the model is forming semantic groupings of drugs based on their active compounds. Optimizing for compound based clustering is similar to optimizing groupings for synonym clustering, thus enabling better discovery of misspellings and synonyms based on cosine similarity in downstream uses of the embeddings.

#### 2.2.3. Final model performance evaluation

We evaluated our word embedding model on the three different word pair similarity tasks. Word pair similarity tasks consist of comparing cosine distances of word pairs within the embedded space with some outside measure of the similarity of those two words. As it is desirable to have an embedded space where distances between word embeddings capture information about the relationship between those two words, with similarity being the relationship of interest for this task. For instance, we would expect a good embedding to embed similar terms like “metastasis” and “tumor” closer to each other in the embedded space than less similar terms like “metastasis” and “finger”.

The first metric is the DBCC, which evaluates the cosine similarity between generic and trademark terms for individual DrugBank drug entries. The second is the UMNSRS Clinical term pair similarity task, where 401 clinical term pairs have human curated similarity ranking [35]. The model similarity ranking was calculated based on the cosine similarity of the vector representations of the terms in a given pair. The third task is the MayoSRS relatedness reference, where human relatedness rankings for 101 clinical term pairs were reported [36]. We evaluated the models based on the Spearman correlation of the human versus model rankings, as for the UMNSRS similarity task.

### 2.3. Identification of candidate tokens

Candidate tokens are tokens in the embedded space that progress to the filtering stage as potential misspellings and slang terms of known terms; we refer to the known term used to generate the list of candidate tokens as the seed term. We identified these based on cosine similarity between the vector representations of known drug terms and embedded tokens. Cosine similarity is not sufficient to define potential candidate tokens as drugs differ in their relative numbers of candidate tokens. For instance, we would expect “Marijuana”, a popular recreational drug, to have more relevant candidate tokens (i.e. misspellings/slang terms) within the text, than a drug like simvastatin, which has no known recreational usage and is rarely mentioned within the corpus. Therefore, we aim to select more candidate tokens for commonly discussed drugs than for rarely mentioned ones. To determine a cosine similarity cutoff for each individual seed term we use the following method. We identified the 1,500 tokens nearest to the seed term within the embedded space, based on cosine similarity. We created noisy labels for these 1,500 tokens by labeling (1) phrase tokens, as discovered in 2.1.2, containing the seed term as positives and (2) labeling other tokens as positives or negatives, based on a term match score. Noisy labels enable us to leverage phrases and tokens that are likely misspellings or synonyms within the data to estimate an appropriate cosine cutoff using standard machine learning metrics, such as an f1-score. Tokens that had a term match score, *f*(*w*_*i*_), greater than 0.4 as positives, a heuristic cutoff that selects an average of 20% of the list as positives based on spot inspections of performance on a few rare and a few high frequency drugs. The remainder of the 1,500 tokens were labeled as negatives. The term match score was calculated as follows:

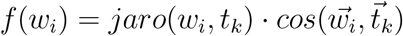

Where *w*_*i*_ is a candidate token contained with the 1,500 nearest tokens and *t*_*k*_ is the seed term used to generate the nearest-neighbor token list. *jaro*(*w*_*i*_, *t*_*k*_) is the Jaro-Winkler similarity between the *w*_*i*_ and *t*_*k*_ tokens, calculated using a prefix weight of 0.1. The Jaro-Winkler similarity is a lexical matching based measure of similarity between two strings. The prefix weight controls the relative impact of matches at the beginning of the strings versus matches made later in the string [37]. 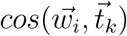 is the cosine similarity between the two tokens’ word vectors. The term match scoring function aimed to balance lexical and semantic similarity of two tokens. The nearest-neighbor cosine similarity cutoff used for pruning for that seed term was then determined by maximizing the f1-score over the noisy labels. Precision and recall values were calculated using the noisy positive and negative labels defined above. Any phrase in the nearest 1,500 tokens that contained an exact match to a known drug term was also included in the set of candidate tokens. This process resulted in a set of candidate tokens, derived from the pruned list of nearest neighbors, and mapped to a seed term, the known drug that served as the clustering centroid. A generic drug was then linked to the union of the clusters for each seed term within that generic drug’s set of associated known drug terms (AKDTs). The set of AKDTs contains all the seed terms for a particular generic drug, based on their DrugBank identifier, for instance the AKDTs for oxycodone would contain oxycodone, oxycontin and oxy ir, as well as the other trademark names for oxycodone formulations. The resulting set of candidate terms for oxycodone would be the union of the candidate token sets of the AKDTs for that generic.

### 2.4. Automated filtering

We compared candidate tokens to their AKDTs. The filtration pipeline is sequential, thus a candidate token that is selected for inclusion in the final term list by an earlier filter is removed from the candidate pool at that stage of the process and not seen by downstream filters. Filtration consists of the following components: comparisons between candidate tokens and their AKDTs using lexical and phonetic metrics (2.4.1, 2.4.2, 2.4.5), lexical comparison to a pill impression database (2.4.3), and the term set intersections between the AKDTs and terms that appear in top 10 results from a Google search for the candidate token (2.4.4).

#### 2.4.1. Edit distance

We computed the edit distance between each candidate token and AKDTs using python-Levenshtein v0.12.0. We included candidate tokens that were within 2 edits of a term in their AKDTs in the final term list.

#### 2.4.2. Partial phonetic and edit distance

We compared the phonetic symbols, derived from the Metaphone algorithm [38], for the first 3 letters of a candidate token to terms in their AKDTs. We then compared candidate tokens with exact matches for the first three phonetic characters to their AKDTs based on normalized edit distance. We included those with a normalized edit distance less than 0.5 in the final term list.

#### 2.4.3. Pill impressions

Pill impressions are the markings made on many pills and tablets by the manufacturer. We compared candidate tokens that were between 1 and 6 characters to pill impressions in the National Drug Codes List. A list of drugs with pill impressions exactly matching the candidate token was intersected with the AKDTs. Candidate tokens with any set intersection were included in the final term list. We found the National Drug Codes List, to be incomplete, thus if an exact pill impression hit was not found in the National Drug Codes List, the drugs.com pill identifier service was queried, and the list of drugs captured in the search list was cross-referenced as noted above and candidate tokens that were a direct match were included in the final term list.

#### 2.4.4. Google search

We registered a Google Custom Search API (https://developers.google.com/custom-search/) and queried candidate tokens through it. We used the top 10 search results to evaluate the candidate token, preprocessing the text content of the title and preview snippets in the same way as the Reddit comment data (2.1.3). We intersected the resulting token set with the AKDTs in the candidate token’s generic group. Candidate tokens with at least one intersecting term were included in the final term list.

#### 2.4.5. Edit distance for low count words

We compared tokens that were captured at the candidate token stage and had less than 30 mentions against all AKDTs sets for lexical similarity. Low count tokens receive special treatment, as they are more difficult to get proper embeddings for, since by definition the corpus contains few contextual examples from which to learn an embedding. Tokens that met the edit distance criteria in 2.4.1 were remapped to the associated seed term and were included in the final term list. In the case of multiple matches, the token was mapped to the AKDT set with the lowest edit distance.

### 2.5. Human rater validation

We gave a random sample of 74 terms derived from 14 seed words there were a subset of 6 generic drug groupings, captured at the candidate term identification stage (2.3), to a group of 12 raters. We presented raters with the associated seed term, the candidate token, and 3 examples of the candidate token being used within Reddit comments. We asked raters to classify the candidate token with a first label, as known (i.e. an exact match to a drug entity in DrugBank), misspelling, synonym, or negative and with a second label as an exact reference to, related to, or unrelated to the associated seed term. For instance, given the seed term “percocet” the first label work as follows: the phrase “taking_percocets” would be labeled as known, “pecocet” would be labeled as a misspelling, “percs” would be a synonym, and “10mg_pills” and “sidewalk” would be labeled as a negative. The second label for “10mg_pills” would be labeled as related, while “sidewalk” would be labeled as unrelated and the remaining examples would be labeled as exact. The related and unrelated classifications were collapsed and designated as non-exact after the rating task was complete. The inter-rater reliability on this 74-term set was used to select a subset of 5 raters with acceptable levels of agreement. Fleiss’ κ was ~ 0.7 for the term type classification task and ~ 0.7 for the relatedness task [39, 40, 41]. These 5 raters individually classified an additional 849 terms derived from 28 seed words that were a subset of 8 generic drug groupings resulting in a total of 923 human annotated candidate tokens. We created the automated labels as follows, edit distance (2.4.1) and the phonetic misspellings (2.4.2) were combined into the “misspelling” category, the pill mark (2.4.3), and Google search (2.4.4) were combined into the “synonyms” category and candidate tokens that were not validated by the automatic methods were designated as “other”. We labeled the human annotated terms that were classified as exact references to the associated seed term as the class they were annotated to and all inexact terms were labeled as “negative”. Performance metrics for the automated classification were calculated by comparing the human and automated labels (Fig. 3).

## 3. Results

### 3.1. Subreddit filtering

The combination of subreddits listed in the r/Health and r/Drugs side bars resulted in 185 subreddits (Supp. File 2) and contained more than 25 million comments. We filtered subreddits based on their medical content enrichment, as described in 2.1.1. The final working corpus was all comments made within the subset of subreddits that had an overall comment enrichment above a cutoff of 1*e*^−4^. This resulted in a corpus of more than 500 million comments over 2,500 subreddits (Supp. File 3).

### 3.2. Word embedding model comparison

We compared the performance of our final Reddit model against 13 publicly available word vector models using various embedding algorithms and corpora [42, 43, 44, 45]. Some words were not represented in the vectorized models (i.e. out-of-vocabulary error), for some of the word pairs in the similarity evaluation task. Nonetheless, we first evaluated all 13 models individually based on the subset of terms that were within the vocabulary of that model (Table 1). We also evaluated the top performing models from the first evaluation based on the subset of term pairs that were within the vocabulary of all those models, to improve interpretation of the results (Table 2). The Reddit model consistently performs best on the DBCC task, while achieving close to top performance on the UMNSRS task (Table 1 and Table 2). Other models outperform the Reddit model on the MayoSRS task.

**Table 1:**
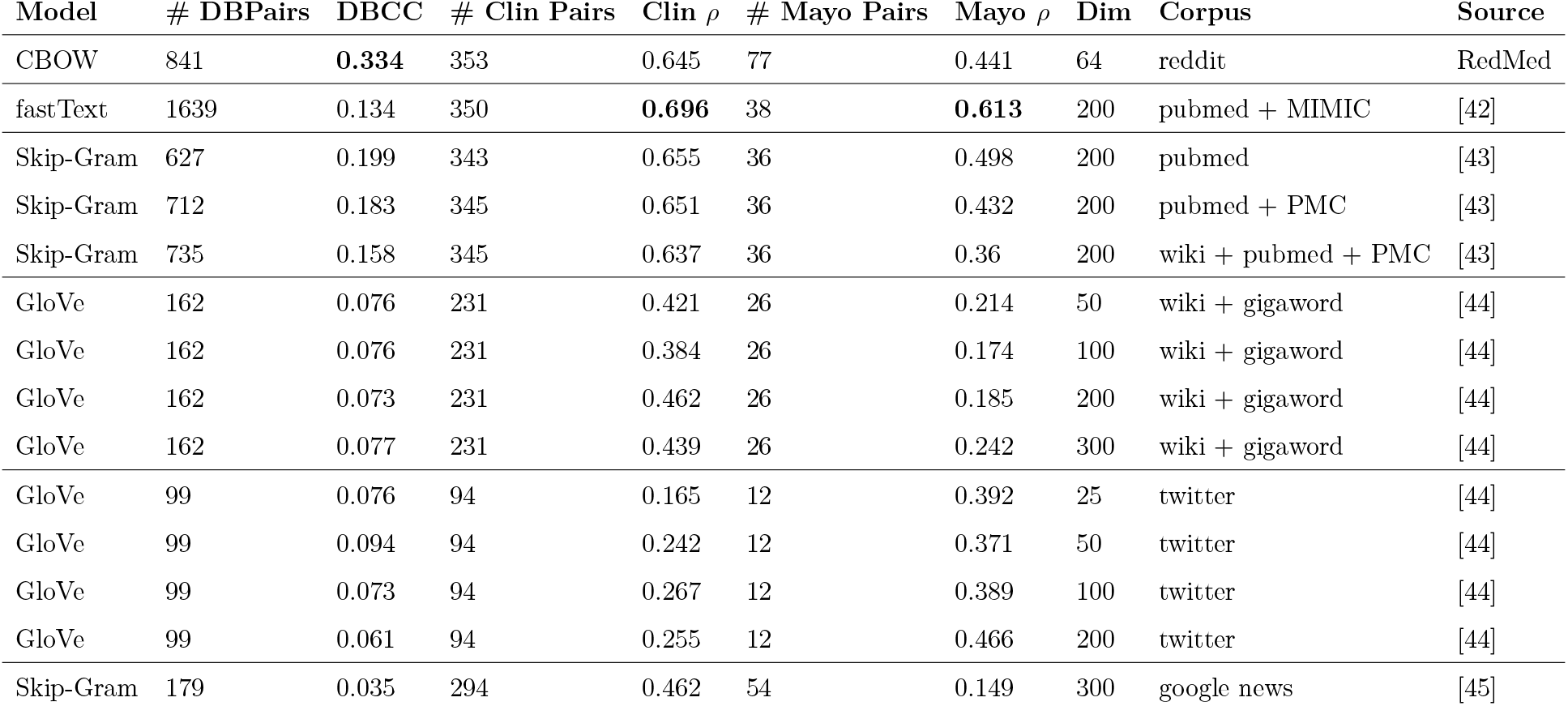
Comparison of Reddit word embedding model to publicly available word embedding models, for per model in-vocabulary term pairs. **#DBPairs** is the number of generic drug groupings, for which there were at least 2 drugs captured by the model. **DBCC** is the DBCC score for that model. **#Clin Pairs** and **#Mayo Pairs** is the number of captured, both terms in the term pair were present in the model, for the UMNSRS Clinical term pairs and MayoSRS term pairs, respectively. **Clin** *ρ* and **Mayo** *ρ* are the Spearman correlations between the model based similarity ranking and the human similarity rankings for the UMNSRS and MayoSRS tasks, respectively. **Dim** is the length of the word vectors for that model. **Corpus** is a brief indication of the corpus(es) used to train the model. **Source** provides a reference for the work from which each model was derived. The Reddit model was the top performing model for the DBCC task. The fastText model trained on PubMed and MIMIC data was the top performing model for the UMNSRS (Clin) and MayoSRS (Mayo) tasks.

| Model | # DBPairs | DBCC | # Clin Pairs | Clin $\rho$ | # Mayo Pairs | Mayo $\rho$ | Dim | Corpus | Source |
| --- | --- | --- | --- | --- | --- | --- | --- | --- | --- |
| CBOW | 841 | **0.334** | 353 | 0.645 | 77 | 0.441 | 64 | reddit | RedMed |
| fastText | 1639 | 0.134 | 350 | **0.696** | 38 | **0.613** | 200 | pubmed + MIMIC | [42] |
| Skip-Gram | 627 | 0.199 | 343 | 0.655 | 36 | 0.498 | 200 | pubmed | [43] |
| Skip-Gram | 712 | 0.183 | 345 | 0.651 | 36 | 0.432 | 200 | pubmed + PMC | [43] |
| Skip-Gram | 735 | 0.158 | 345 | 0.637 | 36 | 0.36 | 200 | wiki + pubmed + PMC | [43] |
| GloVe | 162 | 0.076 | 231 | 0.421 | 26 | 0.214 | 50 | wiki + gigaword | [44] |
| GloVe | 162 | 0.076 | 231 | 0.384 | 26 | 0.174 | 100 | wiki + gigaword | [44] |
| GloVe | 162 | 0.073 | 231 | 0.462 | 26 | 0.185 | 200 | wiki + gigaword | [44] |
| GloVe | 162 | 0.077 | 231 | 0.439 | 26 | 0.242 | 300 | wiki + gigaword | [44] |
| GloVe | 99 | 0.076 | 94 | 0.165 | 12 | 0.392 | 25 | twitter | [44] |
| GloVe | 99 | 0.094 | 94 | 0.242 | 12 | 0.371 | 50 | twitter | [44] |
| GloVe | 99 | 0.073 | 94 | 0.267 | 12 | 0.389 | 100 | twitter | [44] |
| GloVe | 99 | 0.061 | 94 | 0.255 | 12 | 0.466 | 200 | twitter | [44] |
| Skip-Gram | 179 | 0.035 | 294 | 0.462 | 54 | 0.149 | 300 | google news | [45] |

**Table 2:** Comparison of Reddit word embedding model to publicly available word embedding models, for a shared subset of term pairs. For a description of columns see Table 1. The Reddit model was the top performing model for the DBCC task, but the overall DBCC model performance was tighter. The fastText model was the top performing model for the Clin (UMNSRS) and Mayo (MayoSRS) tasks.

| Model | # DBPairs | DBCC | # Clin Pairs | Clin $\rho$ | # Mayo Pairs | Mayo $\rho$ | Dim | Corpus | Source |
| --- | --- | --- | --- | --- | --- | --- | --- | --- | --- |
| CBOW | 279 | **0.181** | 327 | 0.643 | 35 | 0.301 | 64 | reddit | RedMed |
| fastText | 279 | 0.179 | 327 | **0.721** | 35 | **0.63** | 200 | pubmed + MIMIC | [42] |
| Skip-Gram | 279 | 0.177 | 327 | 0.673 | 35 | 0.504 | 200 | pubmed | [43] |
| Skip-Gram | 279 | 0.176 | 327 | 0.655 | 35 | 0.38 | 200 | wiki + pubmed + PMC | [43] |
| Skip-Gram | 279 | 0.162 | 327 | 0.667 | 35 | 0.444 | 200 | pubmed + PMC | [43] |

To evaluate the drug term embeddings within the Reddit model we utilized T-SNE [46] to generate a low dimensional representation of a subset of drugs that have a code designation within the Anatomical Therapeutic Chemical (ATC) Classification System (Figure 2). We visually observe some clustering of sex hormones (G03), antineoplastic agents (L01), and analgesics (N02). The distribution of psycholeptics (N05) and psychoanaleptics (N06) drug classes are broadly overlapping.

### 3.3. Automated validation

Table 3 shows examples of the automated classification categories. We evaluated the classifications made by the automated filtering pipeline against human annotations (Figure 3). The classification accuracy was 0.88 across the 4 classes, with specificity of greater than or equal to 0.90 across all term classes (Table 4). Sensitivity was high (>0.85) for the known, misspelling, and negative classifications, but low for the synonym class.

**Table 3:** Examples of discovered terms and phrases with cosine similarity to the generic term indicated. **Phrases** consists of tokens that contained exact matches to known trademark or generic drug names, and **misspellings** are errors in the spelling of those terms. **Pill marks** were short tokens that matched an entry within a pill impression database. **Synonyms** are candidate tokens whose Google search results contained the associated known drug. Discovered phrases capture relevant information about the drug of interest, such as drug abuse, variation of drug forms (i.e. dosage, strains), and desired attributes (i.e. purity). The misspellings capture both phonetic and lexical errors. Drugs that are primarily delivered in a pill form were sometimes referenced by their pill impressions. Synonym based references to drugs that contain limited lexical similarity to the trademark and generic drug names would likely be missed without semantic information.

| generic | phrases | misspelling | pill marks | synonym |
| --- | --- | --- | --- | --- |
| alprazolam | xanax_addiction (0.59) | zanax (0.62) | g3722 (0.69) | green_hulks (0.71) |
| oxycodone | 80mg_oxy (0.84) | percaset (0.75) | m523 (0.69) | 30mg_blues (0.81) |
| marijuana | cannabis_indica (0.59) | cannibis (0.85) | - | indica_heavy_strain (0.52) |
| cocaine | buy_pure_cocaine (0.28) | cocane (0.71) | - | c17h21no4 (0.67) |

**Table 4:** Sensitivity and specificity for individual term classes. All four classes had high specificity and the known, misspelling, and negative classes had high sensitivity.

|  |  | Automated |  |  |  | Class | Sens. | Spec. |
| --- | --- | --- | --- | --- | --- | --- | --- | --- |
|  |  | known | misspelling | synonym | negative |  |  |  |
| Human | known | 0.86<br>(48) | 0.00<br>(0) | 0.09<br>(5) | 0.05<br>(3) | known | 0.86 | 0.98 |
|  | misspelling | 0.03<br>(3) | 0.92<br>(84) | 0.02<br>(2) | 0.02<br>(2) | misspelling | 0.92 | 0.96 |
|  | synonym | 0.20<br>(9) | 0.13<br>(6) | 0.37<br>(17) | 0.30<br>(14) | synonym | 0.37 | 0.95 |
|  | negative | 0.01<br>(6) | 0.03<br>(24) | 0.05<br>(40) | 0.90<br>(658) | negative | 0.90 | 0.90 |

### 3.4. Proportions of term mentions by class

We searched all publicly available Reddit comments, from 2005-2018, and counted the number of mentions of each term in the discovered term sets. We calculated the proportion of terms for each drug that belonged to each of the 4 classification categories. A subset of drugs and their term proportion breakdowns are presented in Figure 4. Out of all mentions, an average of 79% (± 0.54%) were known, 14% (± 0.47%) were misspellings, 6.4% (± 0.34%) were synonyms, and 0.13% (±0.054%) were pill marks.

## 4. Discussion

In this work, we present both a word embedding model trained on a medically enriched subset of Reddit comments and a set of drug misspellings and slang terms derived from that model. We leveraged edit-distance metrics, existing drug databases, and internet searches to filter candidate tokens in a sequential process (Figure 1).

**Figure 1:**
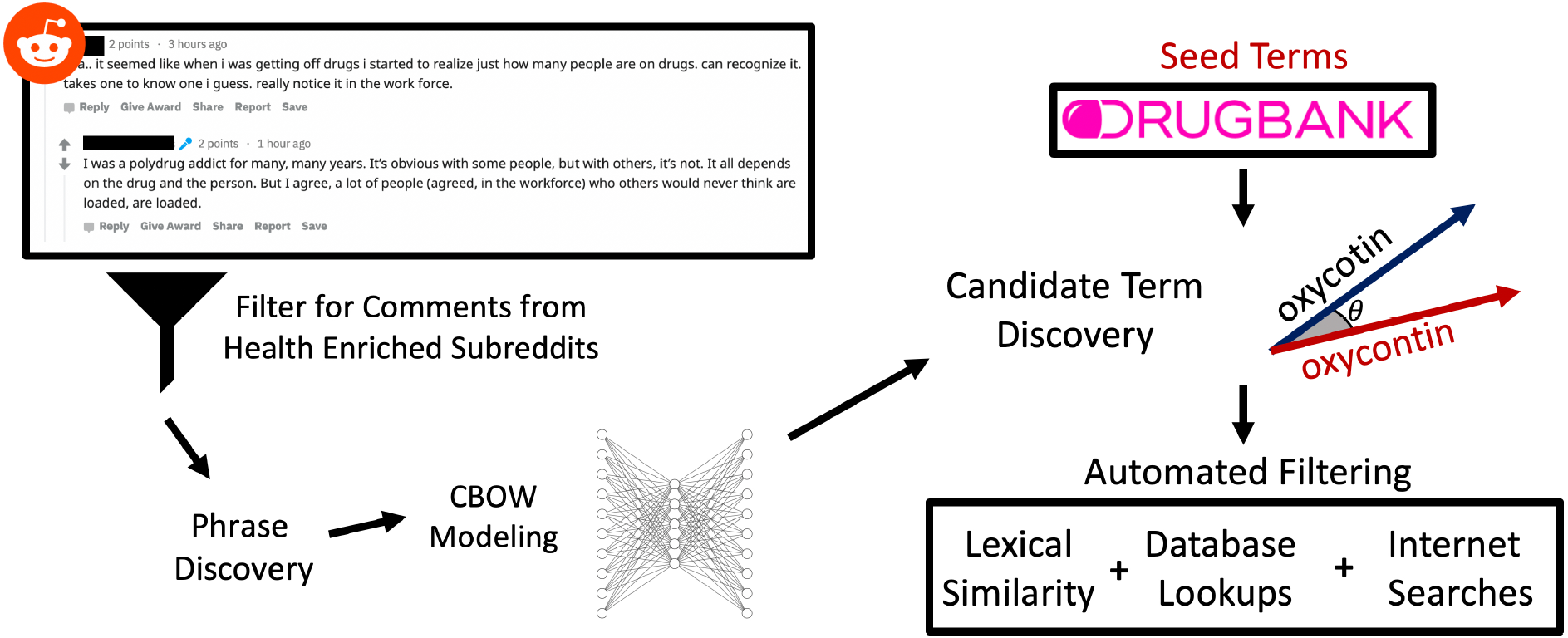
Overview of embedding pipeline. Reddit comments are first filtered based on subreddit for those that contain health related content. The resulting corpus is used for phrase discovery, preprocessed, and used to train a word embedding model. Seed drug terms from DrugBank are used to identify candidate tokens based on their cosine distance to these seed terms. Candidate tokens are then passed through a series of automated filters to yield the final term set.

Although there are several published word embedding models, there were a few reasons we needed to train our own. First, many publicly available word-embedding models were trained on expert generated corpora, such as PubMed, where the underlying semantics, vocabulary, and syntax would be quite different from that found in social media [47]. So while these models might be able to identify synonyms and even misspellings in the expert domain, these are expert synonyms and misspellings not those of the general public. We used Reddit comments to ensure that we captured misspellings and synonym terms, such as slang, that are likely to be unique to social media data. Second, the process through which the data was prepared for these models tries to limit the vocabulary size through preprocessing steps that would have explicitly removed the types of lexical variation we were interested in, such as misspellings. Even if the preprocessing steps for these models did not explicitly exclude or correct misspellings, many sources of expert text undergo multiple rounds of revisions that aim to remove these types of errors from the text. Third, only unigrams or subwords were embedded, while this works for many downstream modeling applications, it would have limited our ability to discover phrases. Phrases are necessary for identifying particular drug synonyms (e.g. “nose_candy”, a slang reference to “cocaine”). Our usage of AutoPhase enabled us to embed these phrases and make them discoverable within the RedMed model. Fourth, models trained on social media data, such as Twitter, that might capture semantic meanings of interest were not enriched for health-related data and thus lacked the majority of drug terms we were interested in.This is primarily due to the rarity of drug mentions within a general social media corpus. Through the selection of specific health-related subreddits we were able to train a social media word embedding model that contains many drugs and other health-related entities of potential interest to the research community.

All three term-pair similarity tasks used to evaluate the word embedding models were defined by medical terminology experts, either database curators or clinicians. Given that our Reddit corpus was derived from comments generated by the general public in an informal setting, we would expect our model to be at a disadvantage for these tasks [47, 48]. The model performance comparison (Table 1 and Table 2) show that despite our model being trained on this layperson corpus, it achieved competitive performance on the UMNSRS task. The RedMed model also outperforms other models on the DBCC task, a somewhat expected result as the DBCC task was used to evaluate hyperparameter settings for our model. Even though other publicly available models outperformed our model on these term-pair similarity tasks, the other publicly available models were poorly suited for our end goal of misspelling and synonym identification. Thus, our model is performing competitively on the similarity-based tasks despite being trained on the Reddit corpus.

We evaluated the RedMed model based on evaluation task labels defined by experts, yet using cosine similarity to a seed term we were able to retrieve relevant candidate terms that include misspellings, slang terms, and pill marks. We find the overall effectiveness of these retrievals to be somewhat surprising. In order for the vector representations to be similar the modeling process requires the words surrounding the known and generic drug names to be similar to the words surrounding the candidate terms within the corpus. For instance the cosine similarity between drug names and their slang terms is quite high, even though we might expect the context surrounding the usage of a proper drug name to differ from the context surrounding a slang term. Users discussing drug abuse might rely more on slang terms and pill marks, with users sharing a pharmacotherapy experience might be more likely to use the proper name for a drug. Our results indicate that there is enough context overlap between these terms for the model to capture that slang terms are indeed references to generic and trademark drugs. On the other hand, we would expect the context to be similar for a drug and many of its misspellings, thus the ability to recover misspellings from the drug name seed words isn’t as surprising and has been used by other groups [18].

The T-SNE visualization offers more evidence of the drug similarities captured by the model (Figure 2). We see tight clustering of particular drug classes, such as antineoplastic agents, sex hormones, and antibacterial agents. We view this as further demonstration that the RedMed model is capturing relevant information about drugs. The distribution of psycholeptics and psychoanaleptics drug classes within the embedded space is overlapping, potentially a result of their shared targeting of psychological phenotypes.

**Figure 2:**
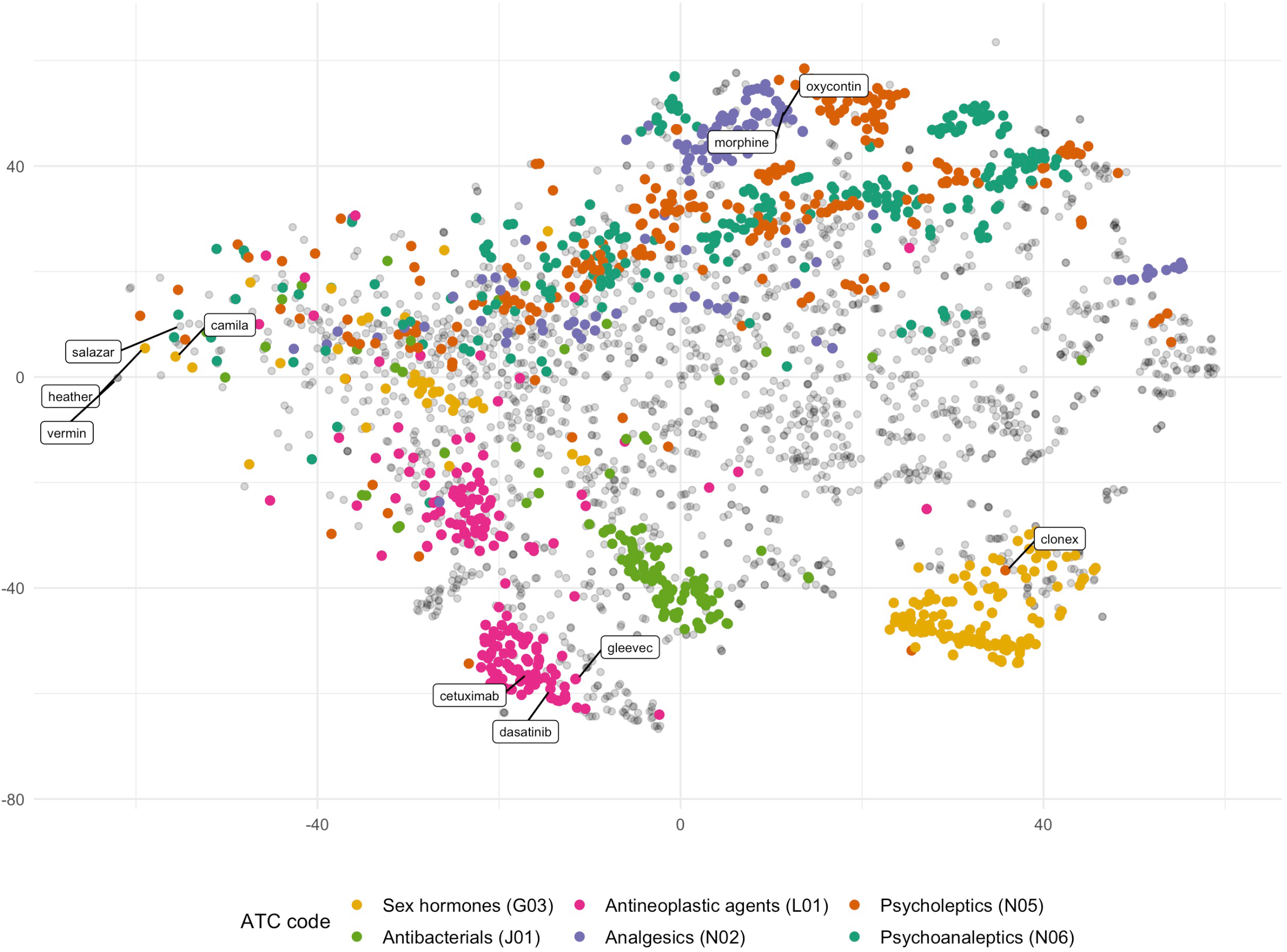
T-SNE visualization of Reddit word embedding space for the subset of drugs with ATC codes. Demonstrates nearness of vectors of drugs within the same ATC drug class in the embedded space. Antineoplastic agents (pink), sex hormones (yellow), and antibacterials (green), some of the larger drug classes, demonstrate closeness within and distance between drug classes in the embedded space. Clustering of drugs within the same therapeutic class indicates that learned representations are capturing drug semantics. Overlapping of psycholeptics (orange) and psychoanaleptics (teal) is a result of the similarity of the therapeutic goal of both drug classes, which is to produce calming effects.

Although, we evaluated the RedMed model on our target task of drug entity identification, exploration of the embedded space yields other concepts that would likely be of interest to public health researchers. For example, looking at terms clustering around “kratom”, a legal opioid receptor family agonist, reveals terms such as “kratom_addiction”, “maeng_da_kratom”, and “previous_opiate_addiction”. The safety of kratom is currently being actively evaluated by the FDA and even this small list of neighboring terms hints at a few potential directions for investigating kratom within the public sphere [49].

Our inclusion of phrases in the embedding model was a critical component of this effort, as phrases were necessary for identifying particular slang, such as “green_hulks”, that could not have been identified from the individual tokens (Figure 3). We allowed misspellings and other unknown English words to remain within the corpus through the preprocessing stage. We were able to leverage both phrases and misspellings as noisy true positives during the candidate term search process (2.3), a crucial component of our pipeline. Additionally, the decision to not limit the corpus to known English words enabled the discovery of methods of referencing drugs we were not expecting, such as pill marks. Given that internet forums contain discussions between individuals at various stages of drug use or abuse, referencing pill marks is a specific means of identifying a particular drug and pill marks allow for drug identification long after pills may have been removed from their original containers.

**Figure 3:**
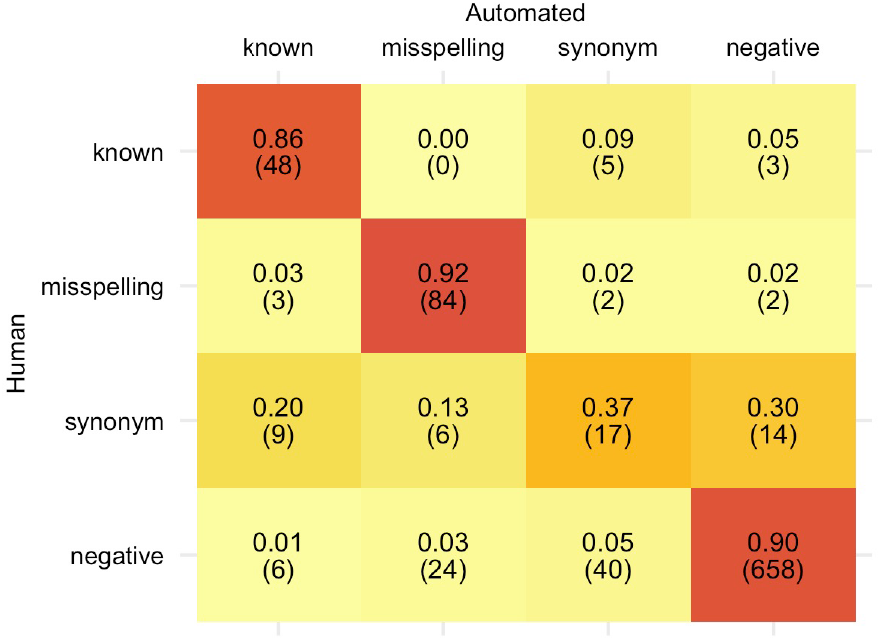
Confusion matrix of human vs. automated validation of candidate token mappings. The five internally consistent human annotators had a high level of agreement with the automated validation pipeline classifications for all term categories except synonyms.

**Figure 4:**
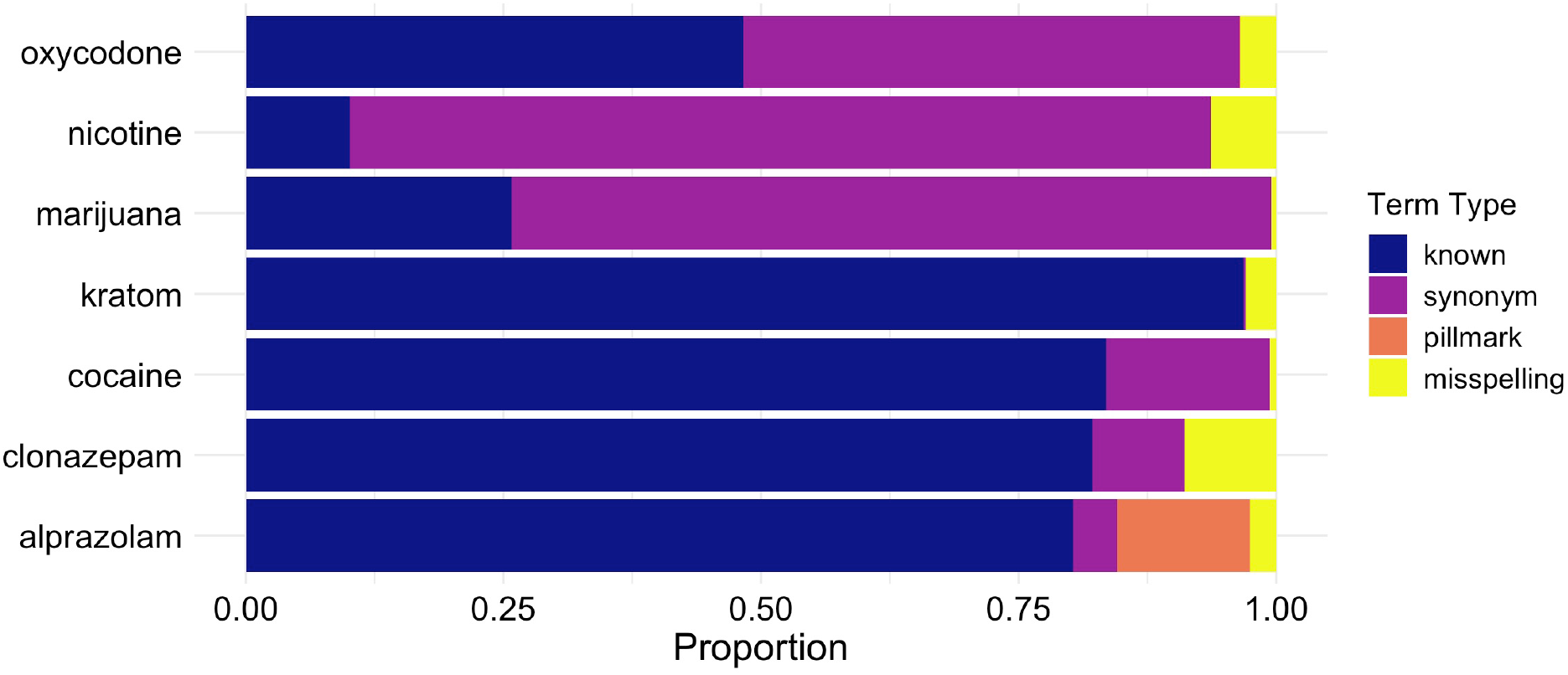
Proportion of all term mentions derived from each of the term types for an exemplary set of seven seed terms. Reddit comment term counts for 2005-2018 were generated for drug terms included in the final term lists. Different term types contribute varying proportions of the overall mentions of a particular drug, with synonyms and misspellings contributing notable proportions of observations. If a search used only known terms as defined by DrugBank drug entries, only the known proportion would be recovered.

The labels we derived from our filtration process achieved high accuracy across the categories for which we saw high human inter-rater agreement, “known”, “misspellings”, and “negative”. For the assignment of a “known” label, our pipeline performs a database lookup and performance primarily relies on the completeness of the database. A misspelling label is assigned based on edit-distance, so primary sources of error are misspellings outside the edit-distance cut-off. We addressed this by allowing a greater edit-distance if the prefix of the word was phonetically identical to the drug to which it was mapped. “negative” was assigned to a term if the above metrics were not satisfied and Google search results did not contain any mentions. We selected Google search as the final validation metric, as we viewed it as mirroring what a human annotator might do for difficult cases. Of course, there are additional nuances in the search results with regards to the specificity of the search term that a human might recognize that our filtration process missed.

Approximately 14% of all drug mentions captured by our term sets were misspellings, this is consistent with a smaller scale study of drug misspellings where Jiang et al. investigated 6 drugs and found around 10% of mentions were misspellings, with higher rates for drugs like codeine and ibuprofen [15]. Misspellings combined with slang terms account for about 20% of the overall mentions. Therefore, social media data collection activities that rely explicitly on exact matches are potentially introducing a large bias into the dataset. For some drugs the problem maybe worse; for instance, while there were 1,812,032 mentions of “marijuana” on Reddit from 2005-2018, there were 5,223,134 mentions of “weed”. While researchers might know to include “weed” in their searches for marijuana posts, other slang terms may not be as obvious to researchers and may represent a large proportion or even a majority of the mentions.

Our ability to identify drug misspellings and synonyms was limited by the overall levels of discussion of particular drugs on the platform. We cannot effectively identify drug misspellings and synonyms for drugs that have few or no mentions within our dataset. This results in relatively few terms for low word count drugs, such as simvastatin. Generally, we still manage to find a few common misspellings and have good coverage of commonly discussed drug.

Some low frequency misspellings and synonym terms were clustered with a different drug than they were actually referencing, resulting in those terms being missed by our approach. For example, “dillaudid”, a misspelling of the trademark drug “dilaudid” clustered close to “oxycodone” in the embedded space, even though the generic form of “dilaudid” is hydro-morphone. We addressed this issue by allowing low count terms, which are more likely to have poor quality embeddings, to be remapped based on our misspelling metrics. Even with this additional step it is unlikely we were unable to fully recover all misspellings. We also note that drug names with alternate meanings outside of the drug context, appear further from their drug classes within the embedded space. The trademark drug names “Salazar” and “Camila” are examples of this effect, as they likely primarily refer to people rather than trademark drugs. This resulted in the vector representations of these drugs that were different than those of other drugs in the embedded space and may have added noise to the clustering process.

Our Google search validation is limited by false negatives when the primary meaning of a candidate token differed from its usage on Reddit. For example, “real_coke”, which refers to real cocaine on Reddit is primarily associated with a soft drink in Google search results. Additionally, Google searches yielded false positives when search results for a relatively general token, such as “painkillers”, returned a reference to the mapped drug in question.

Our term counts were calculated across all comments in the comment archive, without considering the context in which the term was mentioned. While the majority of tokens were not polysemous, certain terms may have suffered from polysemy and thus potentially increased counts with irrelevant mentions. Additionally, our lexicon based approach does not incorporate any information about context and is primarily intended as a means of retrieving data in a keyword search research setting and/or as a means of distant supervision for training more context aware named entity recognition models.

## 5. Conclusions

We present the RedMed model and lexicon, providing an empirical study of drug misspellings and synonym usage in Reddit comment data. We trained the RedMed word embedding model specifically for the task of investigating health-related entities in social media. Although our model is trained on non-technical documents, it performs well on several benchmark medical similarity tasks. We used our method to identify drug misspellings and synonyms. Using the resulting term sets we derived empirical estimates for the rate of misspellings and synonyms usage based on ~3,000 drugs. The RedMed lexicon provides an improved set of search terms for drug-related analysis of social media data (Supp. File 4). Our RedMed model and lexicon are available on Zenodo (https://zenodo.org/record/3238718). Additionally, a python package for text analysis in accordance with our lexicon is available at https://github.com/alavertu/redmed.git.

## Declaration of Competing Interest

The authors have no conflicts of interest.

## Acknowledgements

We’d like to thank the editorial staff and the reviewers for their excellent feedback during the publishing process. This work was supported by the National Science Foundation [DGE – 1656518], National Institutes of Health [R24GM61374, R24GM115264, GM102365, LM05652] and the Food and Drug Administration [U01FD004979].

## Notes

#### Summary of Updates

Additional details have been added to the introduction, methods, and discussion sections. Figure 1 has been revised.

https://zenodo.org/record/3238718

https://github.com/alavertu/redmed.git

